# The paradox of obligate sex: the roles of sexual conflict and mate scarcity in transitions to facultative and obligate asexuality

**DOI:** 10.1101/146076

**Authors:** Nathan W Burke, Russell Bonduriansky

## Abstract

Recent theory suggests that male coercion could contribute to the maintenance of obligate sex. However, it is unclear how sexually antagonistic coevolution might interact with mate scarcity to influence the probability of invasions of obligately sexual populations by mutants capable of parthenogenetic reproduction. Furthermore, if invasion does occur, it is unclear which factors promote or prevent the complete loss of sex. Using individual-based models, we show that male coercion cannot prevent the invasion of a mutant allele that gives virgin females the ability to reproduce parthenogenetically because mutants always benefit by producing at least some offspring asexually prior to mating. Indeed, the likelihood of invasion generally increases as sexual conflict intensifies, and the effects of sexual conflict and mate scarcity can interact in complex ways to promote invasion. Nonetheless, we find that coercion prevents the complete loss of sex unless linkage disequilibrium can build up between the mutant allele and alleles for effective female resistance. Our findings clarify how costs and limitations of female resistance can promote the maintenance of sexual reproduction, turning sex into an evolutionary trap. At the same time, our results highlight the need to explain why facultative reproductive strategies so rarely evolve in nature.

## INTRODUCTION

The prevalence of costly obligate sexual reproduction in some groups of organisms, especially animals, represents a paradox (Burke and Bonduriansky 2017). Theory suggests that facultative strategies that incorporate both sexual and asexual reproduction provide all the genetic advantages of obligate sex but with much lower costs (Green and Noakes 1995; D’Souza and Michiels 2010). For example, facultative sex/asex is as effective as obligate sex at enhancing purifying selection (Lynch and Gabriel 1983; Wagner and Gabriel 1990), creating advantageous allele combinations (Kondrashov 1984; Bell 1988; Hurst and Peck 1996), promoting adaptation (Lynch and Gabriel 1983; Sasaki and Iwasa 1987), and facilitating evolutionary escape from coevolving parasites (Yamauchi 1999; Flatt et al. 2001; Yamauchi and Kamite 2003). Thus, to explain the paradox of obligate sex, theory must account for the capacity of obligately sexual populations to resist invasions by facultatively asexual mutants (Burke and Bonduriansky 2017).

The widespread occurrence of obligate sexuality suggests the existence of mechanisms or dynamics that act as persistent impediments to evolutionary transitions to facultative or obligate asexuality in diverse sexual lineages. One possible mechanism that could impede such transitions is coercion by males (Kawatsu 2013a,b, 2015; Burke et al. 2015; Gerber and Kokko 2016; Burke and Bonduriansky 2017). Males typically benefit from each additional mating whereas females have a lower optimum mating rate, and selection therefore favours male strategies that coerce females into mating even if this results in reduced female fecundity or longevity (Parker 1979; Martin and Hosken 2003; Arnqvist and Rowe 2005; Maklakov et al. 2005). In a sexual population, a mutant allele that makes parthenogenetic reproduction possible may be expected to flourish due to the demographic advantage of producing all-female offspring (Maynard Smith 1978), as well as the ecological and physiological advantages of reproduction without costs of mating. However, coercion could directly inhibit the spread of parthenogenesis by forcing facultative mutants to reproduce sexually, since in many facultatively asexual diploid animals only virgin females are able to reproduce parthenogenetically (Bell 1982). Parthenogenesis may also fail to spread if facultative mutants lose more fitness than sexual females after encountering coercive males. For example, sperm or other male factors could interfere with the proper development of parthenogenetic eggs or offspring (Burke and Bonduriansky n.d.; Schartl et al. 1997; Kawatsu 2013b), or mating could result in increased mortality or reduced fecundity in parthenogenetically reproducing females (Burke et al. 2015). However, male coercion can select for female resistance, potentially setting off a sexual “arms race” (Rice and Holland 1997; Holland and Rice 1998; Gavrilets 2000). Thus, females’ capacity to coevolve effective resistance could play an important role in counteracting the suppressive effect of coercion on parthenogenetic reproduction (Kawatsu 2013a; Burke et al. 2015; Gerber and Kokko 2016; Burke and Bonduriansky 2017).

Conditions that lead to mate scarcity, such as small population sizes or skewed sex ratios, are also thought to be important in the evolution of facultative parthenogenesis in a number of animal groups – including some phasmatids (Schwander and Crespi 2009), ephemeropterids (Brittain 1982), and dipterans (Markow 2013) – because asexual reproduction provides reproductive assurance when females fail to find a mate (Gerritsen 1980; Johnson 1994). The likelihood of encountering mates therefore has the potential to interact with sexually antagonistic selection to either promote or hinder parthenogenesis. A recent theoretical analysis showed that female resistance to mating is more effective at promoting high incidences of parthenogenetic reproduction if population densities are low (Gerber and Kokko 2016); while another theoretical study suggests that obligate parthenogenesis is more likely to evolve if facultative mutants can acquire high resistance (Kawatsu 2013a). However, no previous study has investigated the roles of sexual conflict and mate scarcity simultaneously in an invasion context.

To fill this gap, we investigate how sexual conflict mediated by male coercion interacts with mate scarcity to influence the probability of successful invasion of obligately sexual populations by facultative mutants. Furthermore, if invasion does occur, we investigate the conditions whereby sex is lost through the extinction of males. To address these questions, we employed a series of individual-based simulation models (IBMs) that varied in intensity of sexual conflict, dynamics of sexual coevolution, ecological conditions, genetic background, and relative fecundity of sexual versus parthenogenetic reproductive strategies.

## METHODS

### Overview of IBMs

Using individual-based simulation models (IBMs) in the program NetLogo (Wilensky 1999), we consider a finite population of diploid organisms with overlapping generations inhabiting a gridded environment of square patches in an essentially spherical world. This explicit spatial structure enabled us to create high and low population densities, which are known to affect evolutionary outcomes in facultative systems (Gerber and Kokko 2016), by setting the number of patches to low (11 x 11) and high (51 x 51), respectively. The low density setting simulates conditions that organisms at high risk of mating failure are likely to experience in wild populations (Greenway et al. 2015), and for which, in principle, facultative parthenogenesis would be strongly beneficial for reproductive assurance (Gerritsen 1980); whereas the high-density setting generates high rates of male-female encounter and therefore reduces likelihood of mate scarcity but promotes sexual conflict.

The sexes in our models experience sexual conflict over mating rate. We assume that any more than one mating is costly for females and multiple matings are beneficial for males (Arnqvist and Rowe 2005). We model coercion and resistance in two ways. First, we define coercion and resistance as discrete traits, each controlled by a diploid autosomal locus with two alleles, *c* and *C*, and *r* and *R*, respectively, with additive, sex-limited effects (see Table 1). These ‘discrete-trait’ models investigate cases where one sex gains the upper hand via sexually antagonistic coevolution. Males can gain the upper hand when the *CC* coercion genotype is set to beat all female resistance genotypes (hereafter, ‘coercion’ models), whereas females can gain the upper hand when the *RR* resistance genotype is set to beat all male coercion genotypes (hereafter, ‘resistance’ models; see Table 1). Sexually antagonistic coevolution occurs via selection on standing genetic variation at those two loci, allowing populations to stabilize at coercion-dominated or resistance-dominated states. Second, in a separate set of models (‘continuous-trait models’), we simulate an escalating arms race where coercion and resistance are treated as continuously distributed values (representing a large mutational target) and allowed to coevolve without limit via selection on both standing and mutation variation (see Supplementary Material).

**Table 1.** **Mating outcomes for discrete-trait models** Male coercion genotypes can mate with any female resistance genotype positioned below them in the table, and resistance genotypes can resist mating attempts from any coercion genotype below them in the table. Thus, in coercion models, *CC* males can mate with any female genotype, *Cc* males can mate with *Rr* or *rr* females, and *cc* males can mate with *rr* females. In resistance models, *CC* males can mate with *Rr* or *rr* females, *Cc* males can mate with *rr* females, and *cc* males cannot mate with any females. Similarly, in coercion models, *RR* females can resist mating attempts from *Cc* and *cc* males, *Rr* females can resist mating attempts from *cc* males, and *rr* females cannot resist any mating attempts. In resistance models, *RR* females can resist mating attempts from all male genotypes, *Rr* females can resist mating attempts from *Cc* and *cc* males, and *rr* females can resist mating attempts from *cc* males.

| Coercion model | Resistance model |
| --- | --- |
| *CC* | *RR* |
| *RR* | *CC* |
| *Cc* | *Rr* |
| *Rr* | *Cc* |
| *cc* | *rr* |
| *rr* | *cc* |

We assume that a single sex-limited autosomal locus with two alleles, *p* and *P*, controls reproductive mode. Wild-type *pp* individuals are capable only of sexual reproduction, while *P* is a dominant mutant allele that allows females to reproduce prior to mating (i.e., via facultative parthenogenesis). The *P* allele is introduced either at time-step 0 before the sexes can coevolve, or at time-step 10,001 after sexually antagonistic selection has altered coercion and resistance allele frequencies. We hereafter refer to discrete-trait models set up this way as incorporating ‘no prior sexual coevolution’ versus ‘prior sexual coevolution’, respectively. The *P* allele arises in a random sample of individuals, with 1% of males and 1% of females becoming *PP*, and 1% of males and 1% of females becoming *Pp*. This frequency of *P* alleles limits the extinction of facultative mutants due to drift.

### Initialization of simulations

Simulations start with 500 females and 500 males randomly distributed across patches, each with a fixed lifespan of 100 time-steps and an age of 0 that increases by 1 every time-step. In discrete-trait models, males and females are both allocated coercion and resistance genotypes according to Hardy-Weinberg probabilities and linkage equilibrium, with an allele frequency of 0.5. The carrying capacity of the environment (i.e., maximum population size) is 2,500.

### Life cycle

During each time-step, individuals perform tasks in four ordered phases: moving, mating, reproducing and dying. In the moving phase, individuals turn to face a new direction between 0 and 90 degrees relative to their current direction which is decided by drawing a random number from a uniform distribution with limits 0 and 90. Individuals then move forward one unit (the length of a patch).

During the mating phase, each male randomly chooses a female in his patch that has not mated in the current time-step and tries to mate with her. Males can make only one attempt at mating per time-step, but females can be courted sequentially within a time-step by more than one male if successive mating attempts in a time-step are unsuccessful. Mating occurs when a male’s coercion genotype beats the resistance genotype of the female (see Table 1).

Resistance is either costly or non-costly for females. When resistance is costly, females incur a 10-time-step reduction to their remaining lifespan every time they successfully resist a mating attempt. Following convention (e.g., (Härdling and Kaitala 2005)), we assume that females store enough sperm from one mating to fertilize all their eggs until the end of their life, and that a female’s last mate sires her subsequent offspring.

Males and females incur sex-specific survival costs of mating, applied as penalties of 0, 5, 10, 15 or 20 time-steps deducted from an individual’s remaining lifespan. When the female mating cost is > 0, mating more than once is costly for females, whereas male fitness increases with each additional mating regardless of the cost. Cost of mating to females therefore reflects the intensity of sexual conflict.

Reproduction is a lottery that occurs every time-step if the current population size is less than the carrying capacity. Each female capable of reproducing is allotted a random number from a uniform distribution between 0 and 1. Previously mated females with a random number < 0.1 reproduce sexually. For virgin females that carry the *P* allele, reproduction probability per time-step is globally set at either 0.1 (as for previously mated females) or 0.05 (representing 50%-reduced parthenogenetic fecundity, reflecting a genetic/physiological constraint on asexuality (Lamb and Willey 1979; Engelstadter 2008)). Females can win the reproductive lottery multiple times, but only produce one offspring per reproductive bout. Mated females produce daughters and sons with equal probability, whereas parthenogenesis results in daughters only. As occurs in many facultatively parthenogenetic taxa (Bell 1982), females that mate reproduce sexually thereafter, even if they carry the *P* allele. Although sexual recombination can provide long-term genetic advantages (Hamilton 1980; Kondrashov 1988; Otto and Barton 1997; Peck and Waxman 2000), we ignore these potential benefits to focus solely on short-term invasion dynamics.

Offspring inherit parental alleles and trait values for reproductive mode, coercion, and resistance. We assume that daughters of unmated mothers are produced via apomixis, the most common mechanism of animal parthenogenesis (Bell 1982), and therefore inherit their mothers’ complete genotype. Sexually produced offspring inherit parental alleles following Mendelian rules of segregation.

Following reproduction, an individual’s survival value, *S*, is determined as:

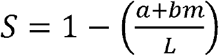

where *a* is an individual’s current age in time-steps, *b* is the sex-specific cost of mating in time-steps, *m* is an individual’s cumulative number of matings, and *L* is the potential lifespan at birth (set at 100 in all models). Death occurs when *S* ≤ 0.

### Analysis

We performed 25 simulation runs for each unique parameter combination of discrete-trait and continuous-trait models to determine the proportion of simulations that ended in *P* allele fixation, *P-p* polymorphism or *P* allele extinction, and the proportion that ended in obligate sex, facultative parthenogenesis, obligate parthenogenesis (male extinction) or population extinction. In one additional run, we collected data every time-step on population size, sex ratio, *P* allele frequency, mean lifetime mating costs, coercion and resistance genotype frequencies, and number of offspring. All simulation runs lasted 20,000 time-steps following the emergence of the *P* allele, except in cases of prior population extinction. Because outcomes for both discrete-trait and continuous-trait models were broadly consistent, we focus below on discrete-trait models, and briefly compare results for those models with outcomes from continuous-trait models. A detailed description of and full results for continuous-trait models are provided in the Supplementary Materials. A list of all model parameters used in discrete-trait models is provided in Table 2.

**Table 2.** **Parameters used in simulations of discrete-trait models**

| Parameter | Description | Parameter levels investigated |
| --- | --- | --- |
| **Number of patches** | The number of patches that make up the world, which determines population density | High (2,601 patches; 'low density')<br>Low (121 patches; 'high density') |
| **Cost per mating for males** | The number of time-steps deducted from male lifespan per mating | 0<br>5<br>10<br>15<br>20 |
| **Cost per mating for females** | The number of time-steps deducted from female lifespan per mating | 0<br>5<br>10<br>15<br>20 |
| **Relative efficacy of coercion and resistance** | The trait that has the upper hand in sexual encounters | 'Coercion can beat resistance' (i.e., males with the most coercive genotype can mate with any female)<br>'Resistance can beat coercion' (i.e., females with the most resistant genotype can resist any mating attempt) |
| **Cost of resistance** | The number of time-steps deducted from the lifespan of females that successfully resist | 0 ('resistance not costly')<br>10 ('resistance costly') |
| **Timing of *P* allele introduction** | The time-step at which mutants carrying the *P* allele are introduced into the population, which determines whether or not populations experience sexual coevolution prior to introduction | 0 ('no prior sexual coevolution')<br>10,000 ('prior sexual coevolution') |
| **Cost of parthenogenesis** | The proportion of fecundity lost by parthenogenetic females relative to sexual females | 0 ('parthenogenesis not costly')<br>0.5 ('parthenogenesis costly') |

Prior to the introduction of the *P* allele in discrete-trait models, relative costs of mating for each sex had consistent demographic effects, with higher costs for one sex generating strongly biased sex ratios (see the first 10,000 time-steps of Figures S1 *A* and *B*). Since large deviations from equal sex ratio are likely to represent extreme cases, we focus below on simulations where male and female costs of mating are balanced, and sex ratios therefore remain approximately equal. We report *P-*allele and reproductive-mode outcomes for all mating cost combinations in Supplementary Materials (Figure S2).

## RESULTS

### Conditions for the invasion of the P allele

We find that the *P* allele spreads via two interacting mechanisms: the ability to reproduce asexually prior to encountering any males (mate scarcity), and the ability to reproduce asexually by resisting males (sexual conflict). The mate scarcity mechanism contributes to the spread of the *P* allele in all versions of the discrete-trait model, including coercion models where males can evolve to coerce any female to mate (Figure 1 *A*). Mate scarcity promotes the spread of the *P* allele in coercion models because at least some virgin females fail to encounter a male by chance, regardless of the effectiveness of male coercion, and the *P* allele gives these females the opportunity to reproduce parthenogenetically. Separate analyses (not shown) confirm that the *P* allele spreads because of this general fecundity advantage and not due to drift. This shows that, under our assumptions, sexual conflict mediated by male coercion cannot impede the invasion of a facultative strategy.

**Figure 1.**
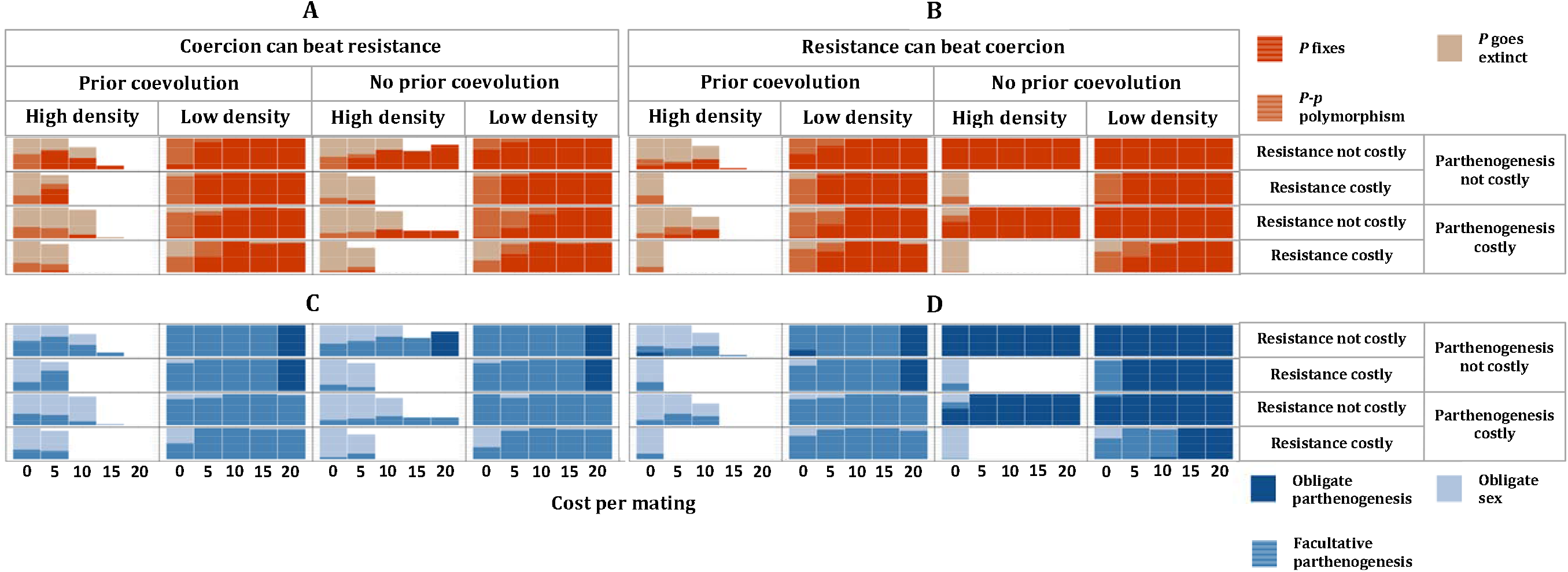
**Evolutionary outcomes following the introduction of the *P* allele for discrete-trait models where the cost per mating is equivalent for each sex**. The cost per mating for females reflects the intensity of sexual conflict, whereas the cost per mating for males is set at the same value as for females to maintain near-equal sex ratios. Left-hand graphs (A and C) are from coercion models; right-hand graphs (B and E) are from resistance models. Upper graphs (A and B) show *P* allele outcomes; lower graphs (C and D) show reproductive mode outcomes. White regions indicate parameter spaces where population extinction occurred before simulations ended. The likelihood of *P* allele fixation (dark orange) increases in panels A and B as costs per mating (and the consequent intensity of sexual conflict) increase along the x-axis. The probability of simulations ending in obligate parthenogenesis (dark blue in panels C and D) also increases as costs per mating increase. The *P* allele easily fixes in panel A, especially when population density is low, because fecundity selection favours the production of parthenogens when mates are scarce, even though males in these simulations can coerce any female to mate. However, the success of the *P* allele in panel A does not translate to widespread obligate parthenogenesis in panel C because coercion ensures the continued production of males when females cannot resist effectively. Instead, obligate sex and facultative parthenogenesis are the most common outcomes in panel C. The *P* allele fixes more often in panel B than in panel A because resistance can beat coercion in these simulations. Transitions to obligate parthenogenesis are also more extensive in panel D than panel C because linkage disequilibrium can build up between the *P* allele and alleles for high resistance when resistance beats coercion and when there is no prior sexual coevolution. However, resistance costs greatly limit the success of parthenogenetic strategies in panels B and D when population density (and therefore sexual conflict) is high. N = 25 simulations for each parameter combination.

In resistance models, positive linkage disequilibrium develops between the *R* allele and the *P* allele, especially when resistance is cost-free, creating strong positive epistasis for female fitness when the capacity for parthenogenesis is coupled with high resistance (Figure 2). When linkage disequilibrium can build up, resistance plays a greater role than mate scarcity in promoting the spread of parthenogenesis during initial stages of invasion (Figure 3 *B*). However, as invasions progress and sexual encounters per female decline with shrinking male sex-ratio as more and more parthenogens are produced, mate scarcity becomes the dominant driver of the *P* allele’s spread (Figure 3 *B*). Likewise, in coercion models with no prior sexual coevolution, resistance partially contributes to the spread of the *P* allele immediately following its introduction because some resistance is still possible at this stage (Figure 3 *A*). However, the *C* allele eventually fixes in these models, and mate scarcity then becomes the sole mechanism by which the *P* allele can spread (Figure 3 *A*). This shows that mate scarcity contributes to *P* allele invasions regardless of whether effective resistance can evolve, but sexual conflict can promote the invasion of facultative mutants if alleles for effective resistance are present in the population and if resisting matings is not too costly.

**Figure 2.**
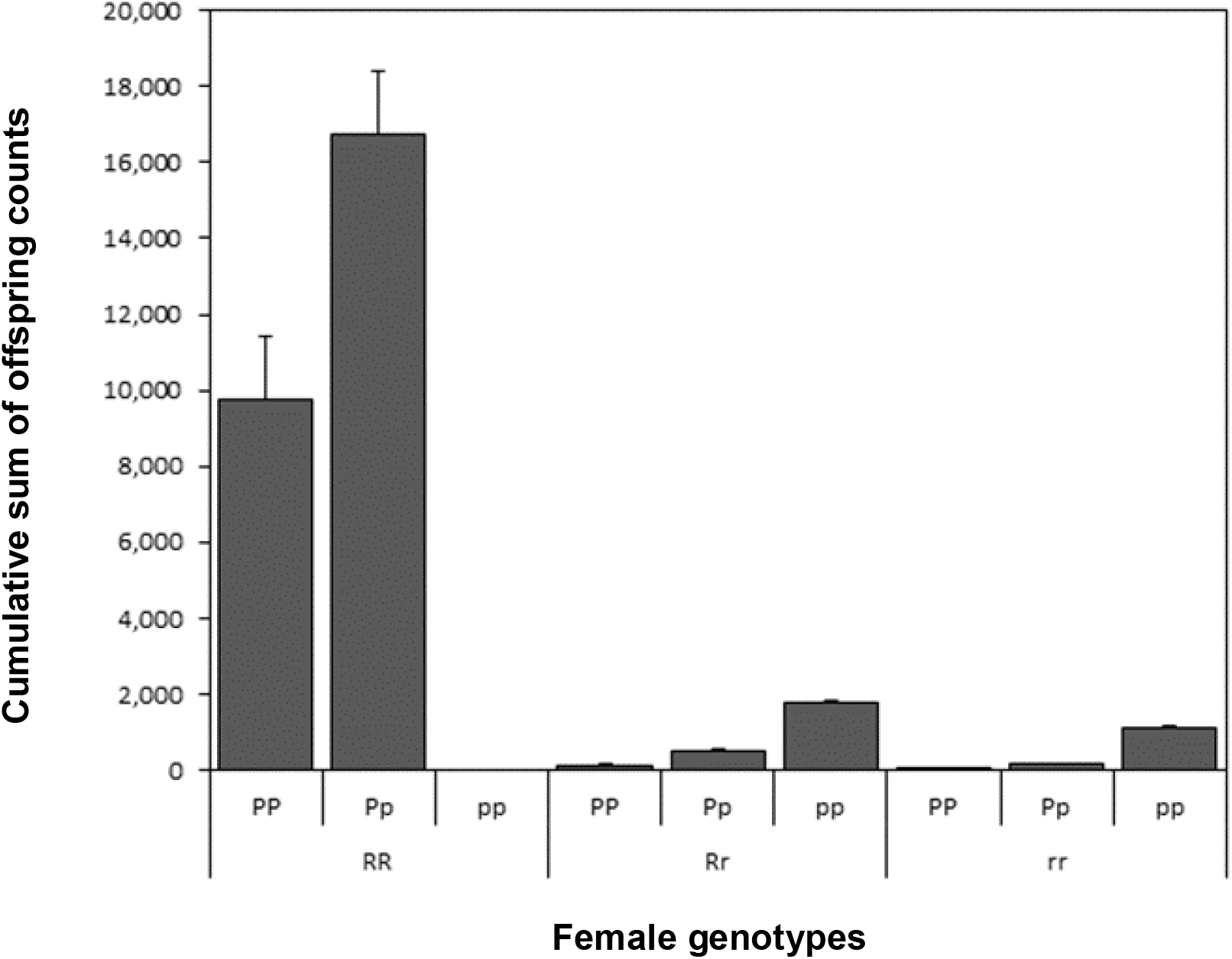
**Mean cumulative sum (+ SE) of offspring produced by female genotypes**. Data are obtained from the first 1,200 time-steps of simulations of resistance models where the *P* allele is introduced without prior sexual coevolution, allowing linkage disequilibrium to build up between *R* and *P*. The parthenogenesis allele *P* is most successful when associated with the most resistant female genotype *RR*. Other parameters are: number of patches: high; cost of resistance: 0; cost of parthenogenesis: 0; female mating cost: 10; male mating cost: 10. N = 25 simulations.

**Figure 3.**
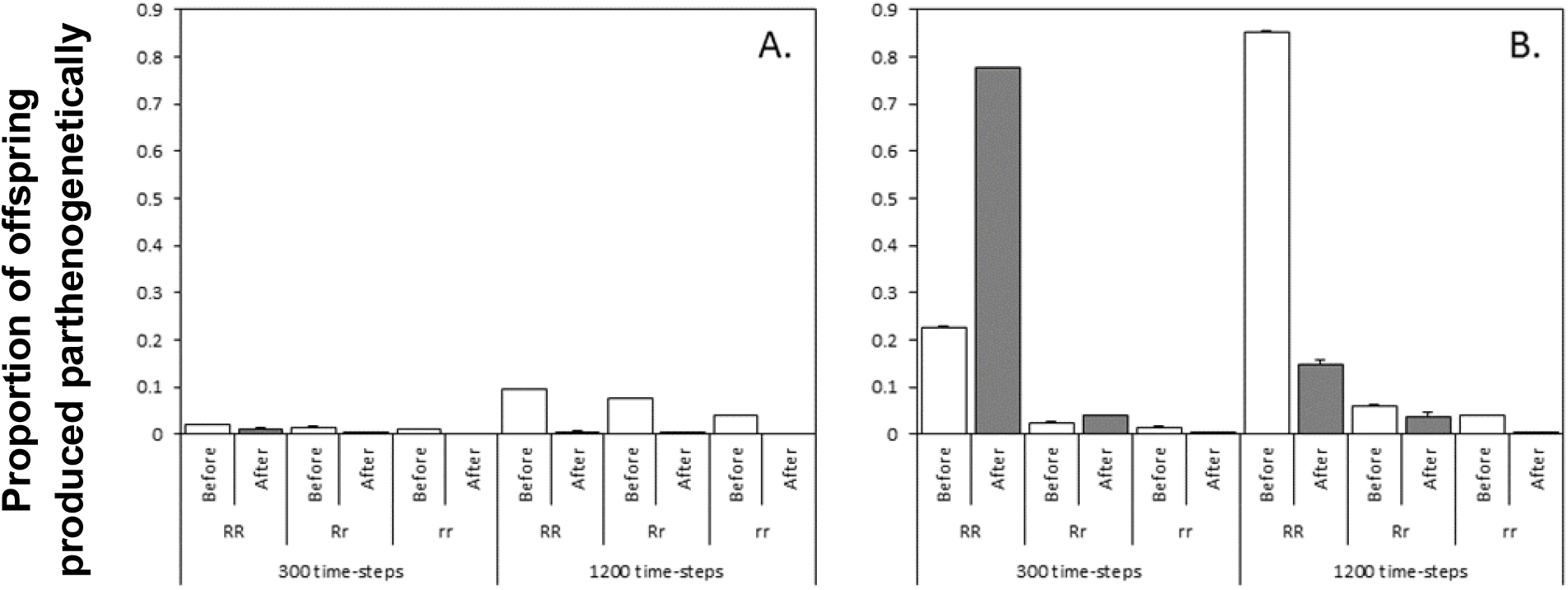
**Mean proportion (+ SE) of offspring produced parthenogenetically by females of different resistance genotypes during the first 300 and 1,200 time-steps of simulation runs before and after initial sexual encounters**. In coercion models (A), most parthenogenetic reproduction occurs before females encounter males (i.e., via the mate scarcity mechanism). This proportion increases as time elapses (contrast the proportion of parthenogenetic reproduction during the first 300 time-steps versus over 1,200 time-steps). Conversely, in resistance models (B), during the first 300 time-steps, parthenogenetic reproduction occurs more frequently *after* mate encounters (i.e., via resistance). But over 1,200 time-steps, parthenogenetic reproduction occurs more frequently *before* mate encounters (i.e., via mate scarcity). In both cases, *RR* females produce the most offspring parthenogenetically. In these simulations, *P* is introduced without prior sexual coevolution, with other parameters as in Figure 2.

The timing of the *P* allele’s introduction also determines whether positive epistasis for fitness between parthenogenesis and resistance develops. This is because the amount of standing genetic variation for antagonistic traits varies depending on whether populations experience sexual coevolution prior to the mutant allele’s introduction. In resistance models with prior coevolution, linkage disequilibrium between the *P* and *R* alleles is unable to build up because females with high resistance fail to mate and reproduce, and selection rapidly eliminates the *R* allele from the population before facultative mutants arise (Figure 4 *B*). However, in resistance models without prior sexual coevolution, genetic variation for resistance is available and thus the *P* allele can rapidly associate with high resistance genotypes and invade over a larger range of the parameter space, especially when resistance is cost-free (Figure 1 *B*). By contrast, timing of introduction has less effect on the *P* allele’s spread in coercion models (Figure 1 *A*) because high coercion rapidly evolves to beat resistance irrespective of prior sexual coevolution (Figure 4 *A* and *C*).

**Figure 4.**
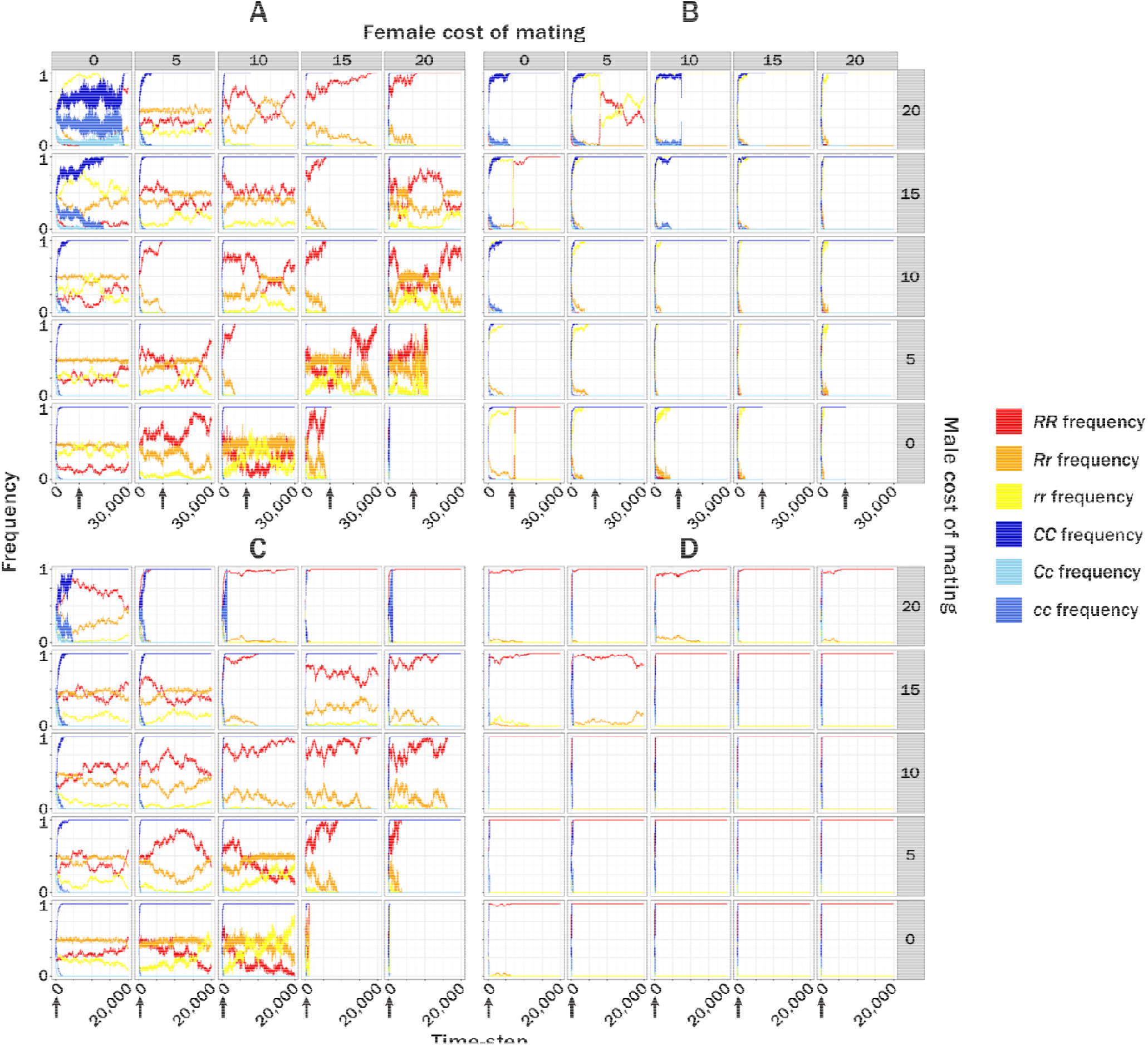
**Time-dependent changes in male coercion and female resistance genotypes for a single simulation per mating-cost combination**. Left-hand graphs (A and C) are from coercion models; right-hand graphs (B and D) are from resistance models. The top graphs (A and B) are from models in which the *P* allele is introduced following prior sexual coevolution; whereas lower graphs (C and D) are from models in which the *P* allele is introduced without prior sexual coevolution. Arrows indicate the time-step at which the *P* allele is introduced. Other parameter settings are: number of patches: high (i.e., low density); cost of resistance: 0; cost of parthenogenesis: 0. In coercion models (A, C), the high-coercion *C* allele (dark blue) rapidly fixes, but directional selection on female resistance ceases following *C* fixation. In resistance models with prior sexual coevolution (B), sexually antagonistic selection fixes both the weakest resistance allele, *r* (yellow), and the strongest coercion allele, *C*; in resistance models with no prior sexual coevolution (D), there is strong directional selection for the high-resistance allele, *R*, as linkage disequilibrium builds up between parthenogenesis and resistance.

The intensity of sexual conflict, reflecting costs of mating for females, also plays an important role in determining the success of *P* allele invasions. When there is no sexual conflict over mating rate (i.e., female cost per mating = 0), extinction of the *P* allele via drift is common (see Figure 1 *A*). However, as the cost per mating increases, sexual conflict enhances the invasion probability of the *P* allele (see Figure 1 *A* and *B*). In high density populations where mating rates per female are very high, female lifespan and opportunities to reproduce decrease with increasing costs of mating, sending populations on a downward spiral to extinction (Figure 1 *A* and *B*). (Resistance costs increase rates of extinction because high density increases the rate of mating attempts and further reduces female lifespan.) These conditions of declining population size are associated with increased rates of *P* allele fixation because the capacity to reproduce without mating is strongly favoured as populations decline and mates become scarce. Together, these results suggest that intense sexual conflict can promote the invasion of alleles for facultative parthenogenesis, thereby potentially averting population extinction.

### Conditions for the establishment of obligate parthenogenesis

We find that the introduction of *P*-allele-carrying mutants into obligately sexual populations leads to one of three distinct evolutionary outcomes: (1) The *P* allele dies out, leaving populations to reproduce via obligate sex; (2) The *P* allele spreads either to an intermediate frequency or to fixation, with males able to persist in the population thereby allowing sex and parthenogenesis to coexist between and/or within individuals (i.e., facultative parthenogenesis); (3) The *P* allele spreads to fixation and parthenogenesis becomes obligate as a result of the complete extinction of males (Figure 2 *C* and *D*). Facultative parthenogenesis and obligate sex are the most common evolutionary outcomes in coercion models because highly coercive males ensure the continued production of male offspring by fertilising eggs of at least some facultatively parthenogenetic females (Figure 1 *C* and *D*). By contrast, obligate and facultative parthenogenesis are the most common results in resistance models. Rapid transitions to obligate parthenogenesis occur across a broad range of parameter space when the *R* and *P* alleles can become linked (i.e., in resistance models without prior sexual coevolution), but only if costs of resistance are not too high (Figure 1 *D*). However, in resistance models with prior sexual coevolution, outcomes for reproductive mode closely resemble those for coercion models (compare Figure 1 *C* and *D*). This is because the high resistance allele *R* (and the potential for linkage disequilibrium) is lost whenever there is prior sexual coevolution (Figure 4 *A* and *B*), and coercion consequently ensures the continued production of males.

### Continuous-trait model

In continuous-trait models where coercion and resistance can escalate in a coevolutionary arms races without limit, we find that *P*-allele frequencies and reproductive mode outcomes are intermediate between those of coercion and resistance discrete-trait models. This occurs because increases in coercion are rapidly counteracted by increases in resistance and vice versa. Thus, linkage disequilibrium between the *P* allele and alleles for high resistance is less likely to result in male extinction because males quickly counter-evolve more effective coercion. Conversely, coercion is less effective at constraining transitions from facultative to obligate parthenogenesis because females quickly counter-evolve more effective resistance which readily becomes linked with the *P* allele. Nonetheless, as with discrete-trait models, *P* allele fixation becomes more likely as the intensity of sexual conflict increases. Detailed results for the continuous-trait model are reported in Supplementary Materials.

## DISCUSSION

Our analysis allowed us to distinguish between instances of parthenogenetic reproduction facilitated by mate scarcity (i.e., parthenogenesis before initial mating attempts) and by resistance (i.e., parthenogenesis after initial mating attempts), and therefore to identify the role of each mechanism in the spread of facultative parthenogenesis and transitions to obligate parthenogenesis. We found that the *P* allele invaded successfully and displaced alleles for obligate sex across most of the realised parameter space due largely to the mechanism of mate scarcity. Even when all matings could be coerced, the *P* allele typically fixed, albeit slowly. High coercion was unable to prevent the invasion of facultative parthenogenesis because fecundity selection favoured females that produced additional offspring prior to encountering a mate, generating positive selection on the *P* allele. However, when successful resistance was possible, the *P* allele invaded across a greater portion of the parameter space because mate scarcity and resistance-mediated mechanisms acted in tandem, making parthenogenesis possible both before and after initial sexual encounters. In other words, the *P* allele conferred the greatest advantage and experienced the strongest positive selection when in positive linkage disequilibrium with alleles conferring a capacity for effective female resistance to mating. High resistance therefore increased the number of offspring produced parthenogenetically, and facilitated rapid and widespread fixation of the mutant allele.

The introduction of the *P* allele into obligately sexual populations led to one of three distinct evolutionary outcomes: obligate sex (*P* allele extinction), facultative parthenogenesis (*P* allele spread), or obligate parthenogenesis (extinction of males and consequent loss of sex). The distribution and frequency of each of these reproductive modes was strongly determined by the genetic architecture of sexual antagonism at the time of the *P* allele’s emergence. When males successfully evolved the capacity to coerce any female to mate prior to introduction of the *P* allele, facultative parthenogenesis and obligate sex were the predominant outcomes, and male extinction rarely occurred. By contrast, when females evolved effective resistance prior to introduction of the *P* allele, male extinction was the most common result. Importantly, this occurred both when coercion and resistance were modelled as traits determined by single loci (and therefore when evolution occurred via selection on standing genetic variation), and when coercion and resistance were modelled as multi-locus traits, representing a large mutational target and allowing sexual arms races to escalate without limit. This suggests that whenever effective resistance cannot evolve, male coercion can impede transitions to obligate parthenogenesis. By contrast, if coercion can be overcome by effective resistance, transitions to obligate asexuality are likely because linkage disequilibrium between the parthenogenesis allele and alleles for effective resistance allows females to avoid mating and its associated costs, leading to the extinction of males.

The limitations of resistance highlighted by our analysis have important implications for understanding the incidence of obligate sex and obligate parthenogenesis in nature. In some species, males appear to “win” sexual arms races due to intense and persistent selection for effective coercion, whether by mechanically overpowering females to force matings (Rowe et al. 1994), by chemical manipulation (Chapman et al. 1995; Andersson et al. 2004), or by pre-copulatory exploitation of sensory biases (Ryan et al. 1993; Holland and Rice 1998). High female resistance genotypes that can resist male coercion may be rare or absent from many populations due to strong selection against absolute resistance, or due to selection favouring convenience polyandry when costs of resistance are high (Rowe 1992). Moreover, many resistance behaviours are plastic, with virgin females often the least resistant to mating ((Ringo 1996); but see (Hosken et al. 2003)), while fixed strategies of high resistance are probably rare in natural populations. The absence of genetic variation for effective resistance may severely inhibit transitions to asexuality by ensuring the continued production of sons. However, parthenogens originating from interspecies hybridisation are often immediately reproductively isolated from their progenitors (Simon et al. 2003), and therefore released from sexual antagonism. This decoupling of parthenogenesis from effective resistance may be one reason why many parthenogenetic animals – including all known obligately parthenogenetic vertebrates (Avise et al. 1992; Simon et al. 2003) – have a hybrid origin.

Theory suggests that facultative strategies should outcompete obligate sex (Green and Noakes 1995; Yamauchi and Kamite 2003). Indeed, our simulations show that facultative parthenogenesis can spread via the mate scarcity mechanism under a broad range of ecological and genetic conditions. Our analysis therefore suggests that the rarity of facultative parthenogenesis in animals may result from the nature of parthenogenetic mutants themselves. First, natural populations probably give rise to facultatively parthenogenetic mutants at very low rates (Schwander et al. 2010) because the complex cytological and physiological changes associated with parthenogenetic reproduction (such as spontaneous development of unreduced eggs) may require simultaneous mutations at multiple loci (Neiman et al. 2014). Second, even when they arise, facultative mutants may be less fecund than wild-type females (Lamb and Willey 1979), especially if the mechanism of parthenogenesis is meiotic (Levitis et al. 2017). For example, facultatively parthenogenetic mutants of the cockroach *Nauphoeta cinerea* produce one tenth as many offspring as non-mutant individuals (Corley and Moore 1999). Our simulations show that even a modest (50%) reduction in fecundity can reduce probability of invasion by parthenogenetic mutants. Third, physiological constraints that prevent females from reproducing parthenogenetically after mating could limit the spread of facultative strategies. Parthenogenetic reproduction after copulation appears to be rare in facultatively asexual diploid animals (e.g., (Chang et al. 2014; Arbuthnott et al. 2015)), except when parthenogenesis is caused by maternally inherited endosymbiont bacteria (Arakaki et al. 2001; Werren et al. 2008). Fourth, the spread of mutants may be further constrained in nature by fitness costs associated with switching from parthenogenetic to sexual reproduction (Burke and Bonduriansky n.d.; Burke et al. 2015). Such genetic constraints on parthenogenetic reproduction could therefore play key roles in preventing the invasion of obligately sexual populations by facultatively parthenogenetic mutants.

Nevertheless, our results generate a number of testable predictions. For example, if male coercion can inhibit the evolution of obligate parthenogenesis, taxa with greater potential for coercion may be less likely to exhibit obligately asexual forms. At a broad phylogenetic scale, the rarity of obligate asexuality in animals compared to plants (Otto and Whitton 2000) may reflect the greater range of opportunities in animal systems for behavioural and chemical coercion, such as chasing or holding mates (Rowe et al. 1994; den Hollander and Gwynne 2009), and transferring toxic ejaculates or anti-aphrodisiacs (Andersson et al. 2004; Wigby and Chapman 2005). Conversely, mate scarcity could play a more important role in plants given their immobility, and this may have selected for facultative asexuality via vegetative reproduction or selfing in many plant lineages. Comparative studies testing these predictions on a finer taxonomic scale may shed light on variation in reproductive strategies within animals, plants, and other eukaryotic lineages. However, such studies will need to quantify actual rates of parthenogenetic reproduction, the incidence and costs of resistance, the costs of mating, and the relative fecundity of sexual versus parthenogenetic reproduction in natural populations of facultative organisms, all of which remain poorly known.

## CONCLUSION

Several recent studies suggest that sexual conflict could play a key role in the maintenance of sexual reproduction, and thus contribute to a resolution of the ‘paradox of sex’ (Kawatsu 2013a,b, 2015; Burke et al. 2015; Gerber and Kokko 2016; Burke and Bonduriansky 2017). However, understanding the role of sexual conflict in the maintenance of obligate sex requires elucidating the ecological, demographic and genetic conditions whereby this factor can promote/inhibit invasions by facultatively asexual mutants in otherwise obligately sexual populations. In particular, given the potential for sexual conflict to interact with population density in facultative systems (Gerber and Kokko 2016), clearly differentiating the role of sexual conflict from the role of mate scarcity in mutant invasions is crucial. In this study, we show that sexual conflict mediated by male coercion cannot prevent facultatively parthenogenetic mutants invading sexual populations because fecundity selection favours mutant females that reproduce prior to encountering males (mate scarcity mechanism). The rarity of facultative parthenogenesis in some lineages therefore suggests important genetic constraints on parthenogenetic reproduction. However, we also show that the probability of facultative populations transitioning to obligate asexuality depends largely on the potential for females to evolve effective, low-cost resistance to mating, and the possibility for linkage disequilibrium to build up between alleles for female resistance and alleles for facultative parthenogenesis. Although females may benefit by reproducing parthenogenetically instead of sexually, obligate parthenogenesis is likely to evolve only if females can overcome male coercion and thereby reproduce without paying the costs of sex. The difficulty of such a feat suggests that sex may be an evolutionary trap imposed on populations by the evolution of coercive males.

## ACKNOWLEDGEMENTS

Thanks to A. Crean, A. Hooper, Z. Wylde, E. Macartney and A. Runagall-McNaull for helpful discussions on model design. We confirm that we have no conflicts of interest in the production of this work.

